# Essential Loop Dynamics Modulates Catalytic Activity in α-Chymotrypsin

**DOI:** 10.1101/2021.08.11.455937

**Authors:** Pritam Biswas, Uttam Pal, Aniruddha Adhikari, Susmita Mondal, Ria Ghosh, Dipanjan Mukherjee, Tanusri Saha-Dasgupta, Sudeshna Shyam Choudhury, Ranjan Das, Samir Kumar Pal

## Abstract

Conformational dynamics of macromolecules including enzymes are essential for their function. The present work reports the role of essential dynamics in alpha-chymotrypsin (CHT) which correlates with its catalytic activity. Detailed optical spectroscopy and classical molecular dynamics (MD) simulation were used to study thermal stability, catalytic activity and dynamical flexibility of the enzyme. The study of the enzyme kinetics reveals an optimum catalytic efficiency at 308K. Polarization gated fluorescence anisotropy with 8-anilino-1-napthelene sulfonate (ANS) have indicated increasing flexibility of the enzyme with an increase in temperature. Examination of the structure of CHT reveal the presence of five loop regions (LRs) around the catalytic S1 pocket. MD simulations have indicated that flexibility increases concurrently with temperature which decreases beyond optimum temperature. Principal component analysis (PCA) of the eigenvectors manifests essential dynamics and gatekeeping role of the five LRs surrounding the catalytic pocket which controls the enzyme activity.

## Introduction

α-Chymotrypsin (CHT) is a proteolytic enzyme of the group S1 serine protease (EC 3.4.21.1) catalyze the hydrolysis of a peptide bond.and are among the most widely distributed enzyme in the biological strata. CHT has attracted considerable interest due to its importance in understanding protein folding or unfolding. CHT play crucial role in several important biological processes such as protein digestion (1, 2), immune response (3, 4) and insect molting (5, 6). CHTs are also known to play a vital role in the intracellular protein turnover (7). CHTs are used extensively in meat industry (8), brewery (9) and in pharmaceutical industries (10, 11).

Studies investigating the role of essential motions in enzyme activity are poorly understood. Previous studies have investigated the correlation between structure and activity. However, most of these have been confined either at the active site or upon binding of small molecules to CHT (12, 13), and provide very little information on the essential motions of enzyme controlling catalysis. Enzyme activity is dependent on the mobility of the protein scaffold and any change associated to such motions can produce different level of selectivity and activity (14, 15). Therefore, it is of our interest to investigate and study the essential motions governing catalysis in CHT.

In the present paper we observe the presence of five loop regions (LRs) near the catalytic S1 pocket of CHT. We hypothesize that these LRs manifest a gatekeeping role controlling catalysis at different temperatures. In the current study, we correlate the essential motions of CHT with its catalytic activity using steady state, circular dichroism (CD) and picosecond resolved spectroscopy along with all atom molecular dynamics (MD) simulation and principal component analysis (PCA).

## Materials and methods

### Materials

Alpha-chymotrypsin (≥ 40 U mg^−1^), Ala-Ala-Phe-7-amido-4-methylcoumarin (Ala-Ala-Phe-AMC), and 8-anilino-1-naphthalenesulfonic acid (ANS) (≥ 97%) were purchased from Sigma (Saint Louis, USA). All solutions were prepared in phosphate buffer (10mM, pH 7.0) without any further purification using water from Milipore system unless stated otherwise.

### Sample preparation

CHT-ANS complex for anisotropy analysis was prepared by mixing ANS (0.5 μM) with CHT (5 μM) in phosphate buffer (pH 7.0) under continuous stirring for 5 h at 4 □, which was dialysed to remove the free probe (16).

### Circular dichroism (CD) spectroscopy

Temperature dependent (278K-338K) circular dichroism (CD) spectroscopy study was conducted with 2 μM CHT solution using a quartz cuvette of 1 mm path length in a Jasco 815 spectrometer with temperature controller attachment. Spectral deconvolution into relevant secondary structure was done using CDNN software (17).

### Enzyme activity assay

Temperature dependent (278K-338K) enzymatic activity studies were conducted using 0.6 μM CHT concentration and the substrate Ala-Ala-Phe-AMC (λ_max_ 325 nm) concentration was varied between 20-200 μM. Substrate was cleaved to produce 7-amido-4-methyl coumarin (λ_max_ 370 nm). The absorbance was monitored at 370 nm using a Shimadzu Model UV-2600 spectrophotometer with an attachment for temperature dependent studies (TCC 240 A). Michaelis-Menten kinetics parameters were calculated using a Lineweaver-Burk plot (18).

### Polarization-gated anisotropy measurements

Time resolved fluorescence transients were measured in a LifeSpec-ps picosecond diode laser-pumped fluorescence spectrophotometer from Edinburgh Instruments (Livingston, UK). The picosecond excitation pulses from the picoquant diode laser were used at 375 nm for the excitation of ANS with an instrument response function (IRF) of 80 ps with an attachment for temperature dependent studies. A microchannel-plate-photomultiplier tube (MCP-PMT; HamamatsuPhotonics, Kyoto, Japan) was used to detect the photoluminescence from the sample after dispersion through a monochromator. The flexibility of CHT was monitored by measuring time resolved fluorescence anisotropy (r(t)) decays of ANS (16, 19, 20) in complex with CHT at different temperatures from 278K to 338K. Time-resolved anisotropy, r(t), was determined from the following equation.

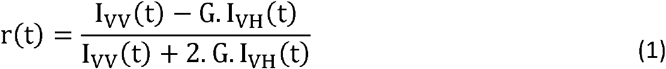

where I_VV_(t) and I_VH_(t) are parallel and perpendicular polarizedfluorescence decays of the dye, respectively, recorded using avertically polarized excitation light. G is an instrument andwavelength dependent correction factor to compensate for the polarization bias of the detection system and its magnitude was obtained by a long tail matching technique (21, 22). The anisotropy decays could be satisfactorily fitted by a multi-exponential decay model

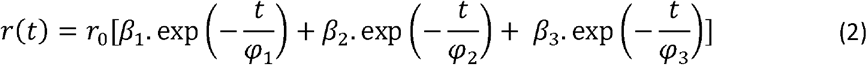

where r_0_ is initial anisotropy, φ_i_s are rotational relaxation times, and β_i_s are their corresponding amplitudes.

### Classical molecular dynamics (MD) simulation

Temperature dependent dynamical behaviour of CHT was further studied by molecular dynamics (MD) simulation. CHT structure (1CGJ) was subjected to energy minimization by Schrodinger Maestro 2018-1 (Academic release). Simulations were carried out at 278K, 288K, 308K, 328K and 338K. CHT was placed in a periodic boundary box at a minimum distance of 10Å from each side. The enzyme was solvated with pre-optimized simple point charged (SPC) water molecules. Then, the charge of the system was neutralized by adding necessary ions (seven Na^+^). The system was equilibrated using a five step relaxation protocol (23). Normal pressure (1 bar) and defined temperature with no particle restraints were used for the final run. A Nose ‘–Hoover thermostat and a Martyna–Tobias–Klein barostat maintained the temperature and pressure of the system, respectively. MD simulation was run in an OPLS 2005 force field (24) with a short range coulombic interaction cut-off of 9 Å. Long range coulombic interactions were handled by a smooth particle mesh Ewald method with a Ewald tolerance of 10^−9^. The simulations were run for 30 ns at the said temperatures. The overall structural changes in the protein with time were computed in terms of the root mean square deviation (RMSD). The radius of gyration (Rg) and residue-wise fluctuations (RMSF) were also computed from the simulation trajectory.

### Normal mode analysis

MD simulation data helps in identifying the configurational space in which anharmonic motion takes place by reduction of dimentionality. The principal component analysis (PCA) takes the trajectory of a molecular dynamics simulation and extracts the dominant modes in the motion of the molecule (25–27). These dominant motions correspond to the correlated vibrational modes or collective motions of the groups of atoms in the normal mode analysis. The principal component analysis (PCA) method depends on the covariance matrix to predict the atomic displacement of protein (28). The direction and amplitude of the dominant motions along a simulation trajectory by principal component analysis (PCA) were computed by Normal Mode Wizard in VMD 1.9.3 program (29) using ProDy (30). The porcupine plots and mobility plots were generated for the respective temperatures by the program displaying a graphical summary of the motions along the trajectory. Supplementary video files of motions were also prepared by VMD (29).

## Result and discussion

### Structural analysis

Thermal stability of CHT was quantified by circular dichroism (CD) spectroscopy in the far UV region. The far UV-CD spectra of CHT was characterized by a minimum at ~202 nm with no positive band (31, 32) (Fig.1a). Deconvolution of the spectra displays insignificant changes in the CHT secondary structure indicating to its thermal stability and absence of widespread thermal denaturation of the enzyme within the temperature range of our study. 1.4% decrease in α-helix, 2.6% decrease in β-sheet, 1.1% increase in random coil were observed (Table.1) (Fig.1b). These are reflective of an insignificant change in the secondary structure of CHT upon thermal treatment, suggesting thermal stability of the enzyme within the limitations of our study. To understand and correlate the effect of temperature on the overall geometry and compactness of the enzyme 30 ns all atom MD simulations were run at different temperatures-278K, 288K, 308K, 318K, 338K. The MD trajectories were investigated for RMSD and radius of gyration (r_gyr_). RMSD is a parameter for assessing the stability of MD trajectories. RMSD signifies the deviation from the initial structure which indicates a change in geometry of the enzyme to the initial structure (15). It was quite evident from Fig.1c that the structure reached an equilibrium geometry after ~20 ns. The RMSD was steady over time at different temperatures and had reached a stable conformation. To understand the effect of temperature on the overall compactness of the enzyme the radius of gyration (r_gyr_) was calculated. rgyr is a measure of the average distribution of atoms from their common center of mass. Lower r_gyr_ values corresponds to a compact structure while higher r_gyr_ describe a loose packing of the structure. As documented from Fig.1d we observe no significant change in the overall compactness of the enzyme during the course of the simulation event corroborating with our circular dichroism (CD) Study.

**Fig.1:**
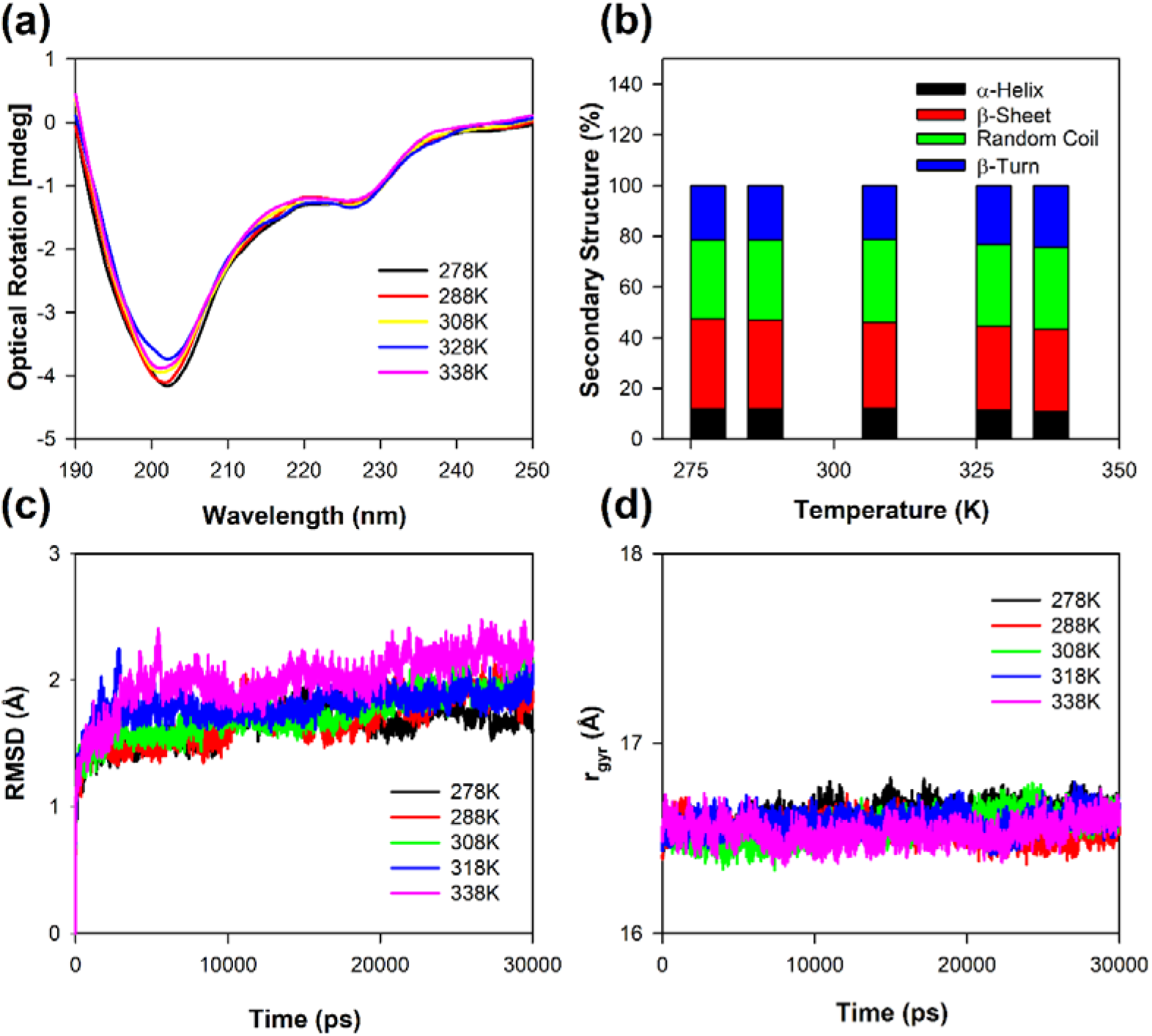
Structural analysis and thermal stability of CHT. (a) Far UV CD spectra of CHT at different temperatures. (b) Secondary structure analysis of CHT. Insiginificant changes to the CHT secondary structure was observed. (c) RMSD analysis of CHT over 30 ns MD simulation event at different temperatures. (d) r_gyr_ of CHT over 30 ns simulation event at different temperatures. Both the RMSD and r_gyr_ analysis indicated no major disruption to the CHT architecture.

**Table.1:** Temperature dependent secondary structure of CHT as deconvoluted by CDNN

| Temperature (K) | % α-Helix | % β-Sheet | % Random Coil | % β-Turn |
| --- | --- | --- | --- | --- |
| 278 | 12 | 35.30 | 31.20 | 21.50 |
| 288 | 12 | 35.10 | 31.60 | 21.30 |
| 308 | 12.10 | 34 | 32.70 | 21.20 |
| 328 | 11.40 | 33.20 | 32.10 | 23.30 |
| 338 | 10.60 | 32.70 | 32.30 | 24.40 |

### Enzymatic activity study

A detailed kinetics of the CHT protease activity was studied in temperatures ranging from 278K to 338K. For different concentrations (20-200 μM) of Ala-Ala-Phe-AMC (substrate) the initial velocity corresponding to protease activity were measured. Lineweaver-Burk plot was used to determine the kinetics parameters (K_m_, V_max_, k_cat_, and k_cat_/K_m_). CHT belongs to the class of serine protease that catalyzes the hydrolysis of Ala-Ala-Phe-AMC (substrate) into 7-amido-4-methycoumarin (Fig.2a & 2b). Fig.2c & 2d illustrate the change in the product formation and catalytic efficiency upon temperature change. In the study thermal treatment was found to enhance catalytic efficiency, reaching maximum efficiency at 308K. Further increase in temperature beyond 308K led to fall in catalytic efficiency. It should also be noted that low catalytic efficiency and turnover was observed at temperatures below 308K. The result is consistent with our thermal stability study indicating the favourable formation of a stable enzyme-substrate complex over a wide range of temperatures. The summary of the kinetics parameters are shown in Table.2.

**Fig.2:**
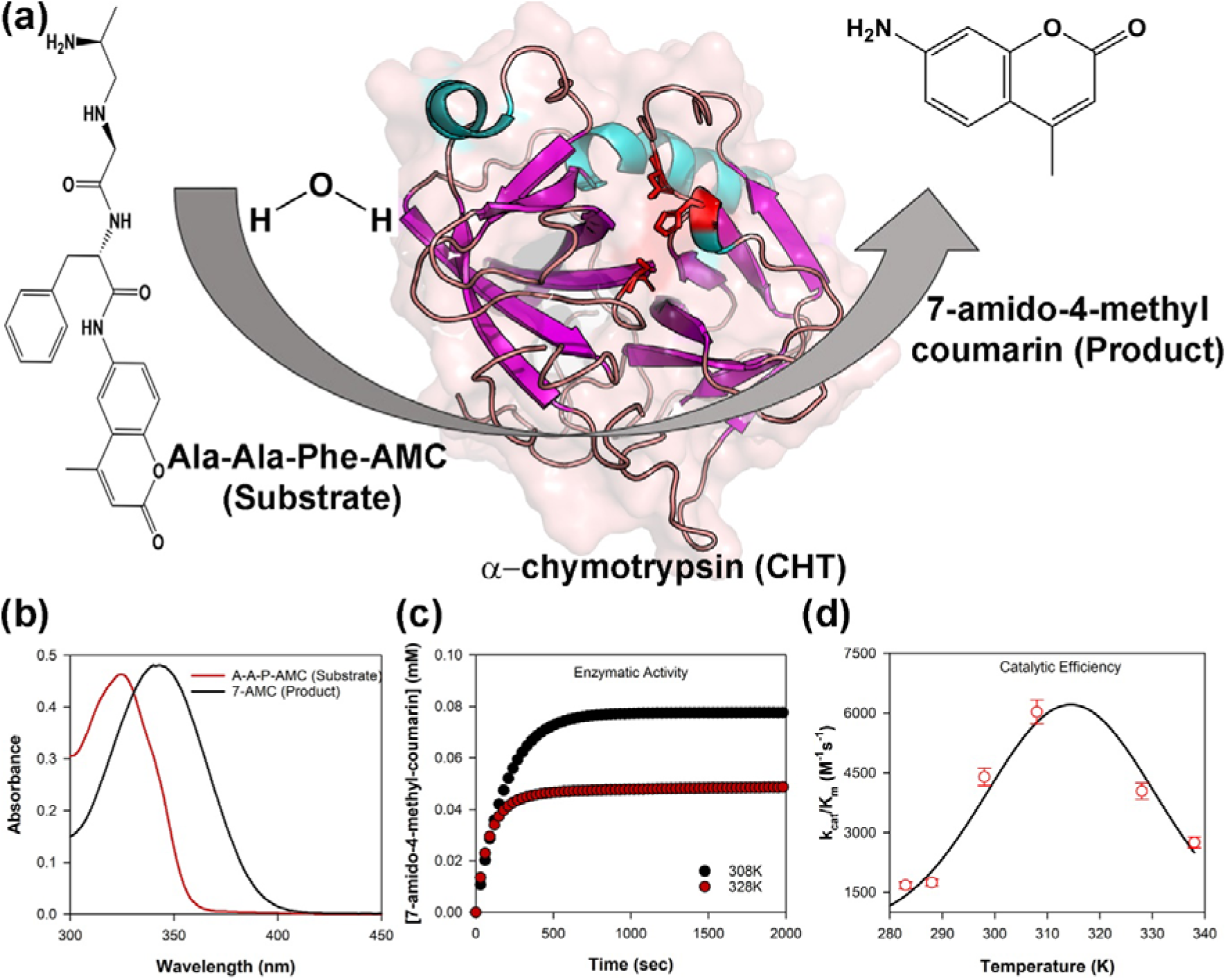
Showing the temperature dependent enzyme catalysis of CHT. (a) Skeletal outline of reaction catalysed by CHT. (b) Absorbance spectra of substrate Ala-Ala-Phe-AMC and the catalysed product 7-amido-4-methyl coumarin. (c) Effect of different representative temperature on product formation. An increase in temperature showed reduced product formation beyond 308K. (d) Catalytic efficiency (k_cat_/K_m_) of CHT at different representative temperature generate a typical bell shaped curve. Optimum temperature at 308K. Beyond 308K there was a decrease in efficiency.

**Table.2:**
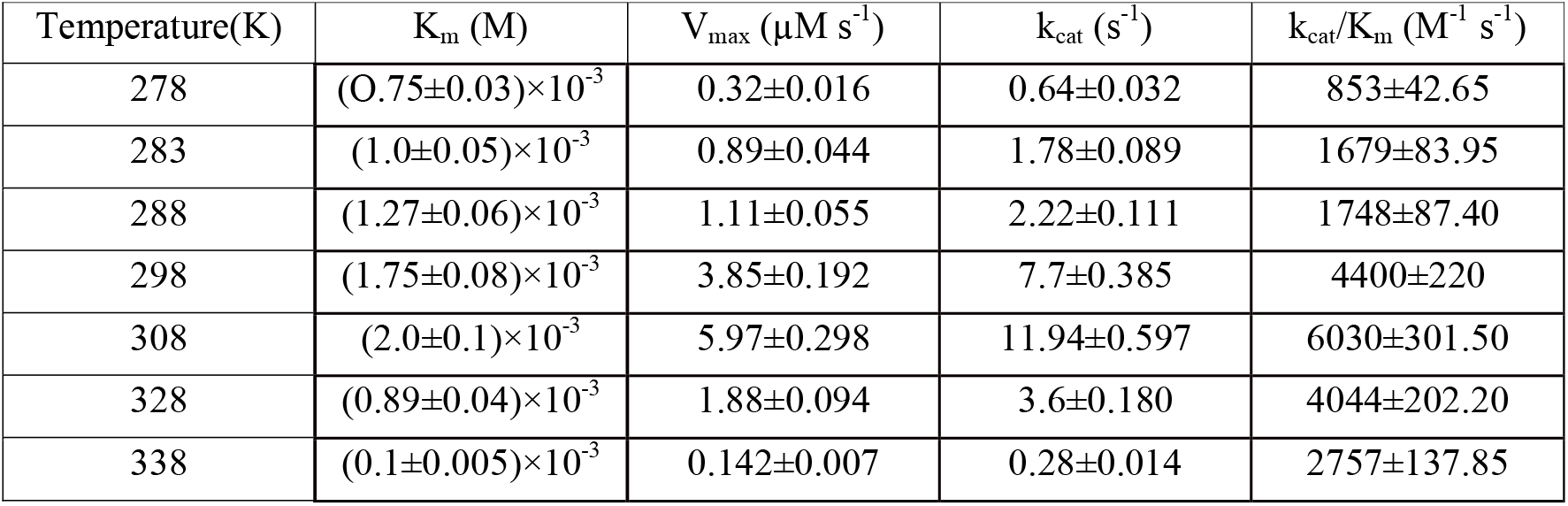
Temperature dependent kinetics of CHT catalysed hydrolysis of Ala-Ala-Phe-AMC

### Polarization-gated fluorescence anisotropy

Time resolved fluorescence anisotropy (r(t), Fig.3) decays of ANS (16, 19, 20) in CHT at different temperatures are characterized by three rotational correlation times (Table 3). The time constant (φ_1_) of ~54-67 ps corresponds to orientational relaxation of the dye in water due to its close resemblance to the rotational relaxation time of ANS in water (~70 ps) (16). On the other hand, the longer sub-nanosecond component (φ_2_) may be ascribed to segmental motions of the amino acid residues Cys-1-122 of the protein, because, ANS binds rigidly at a single site on the surface of CHT near the Cys-1-122 disulfide bond (16) which is localized almost opposite to the catalytic centre of the enzyme. The longest time constant (φ_3_) represents the global tumbling motion of the ANS/CHT complex. Upon an increase of temperature the sub-nanosecond component (φ_2_) remains more or less unchanged, until at 308K it decreases significantly to 550 ps (Table 3). This is reflective of significantly increased segmental mobility of the amino acid residues opposite to the catalytic centre. As a consequence of such increased flexibility of the amino acid residues opposite to the catalytic centre it likely becomes increasingly accessible to the substrate resulting in optimal enzymatic activity of CHT. On further increase of temperature the rotational relaxation time (φ_2_) increases indicating decreased flexibility of the amino acid residues Cys-1-122 with a consequent decrease in the catalytic efficiency of the enzyme.

**Fig.3:**
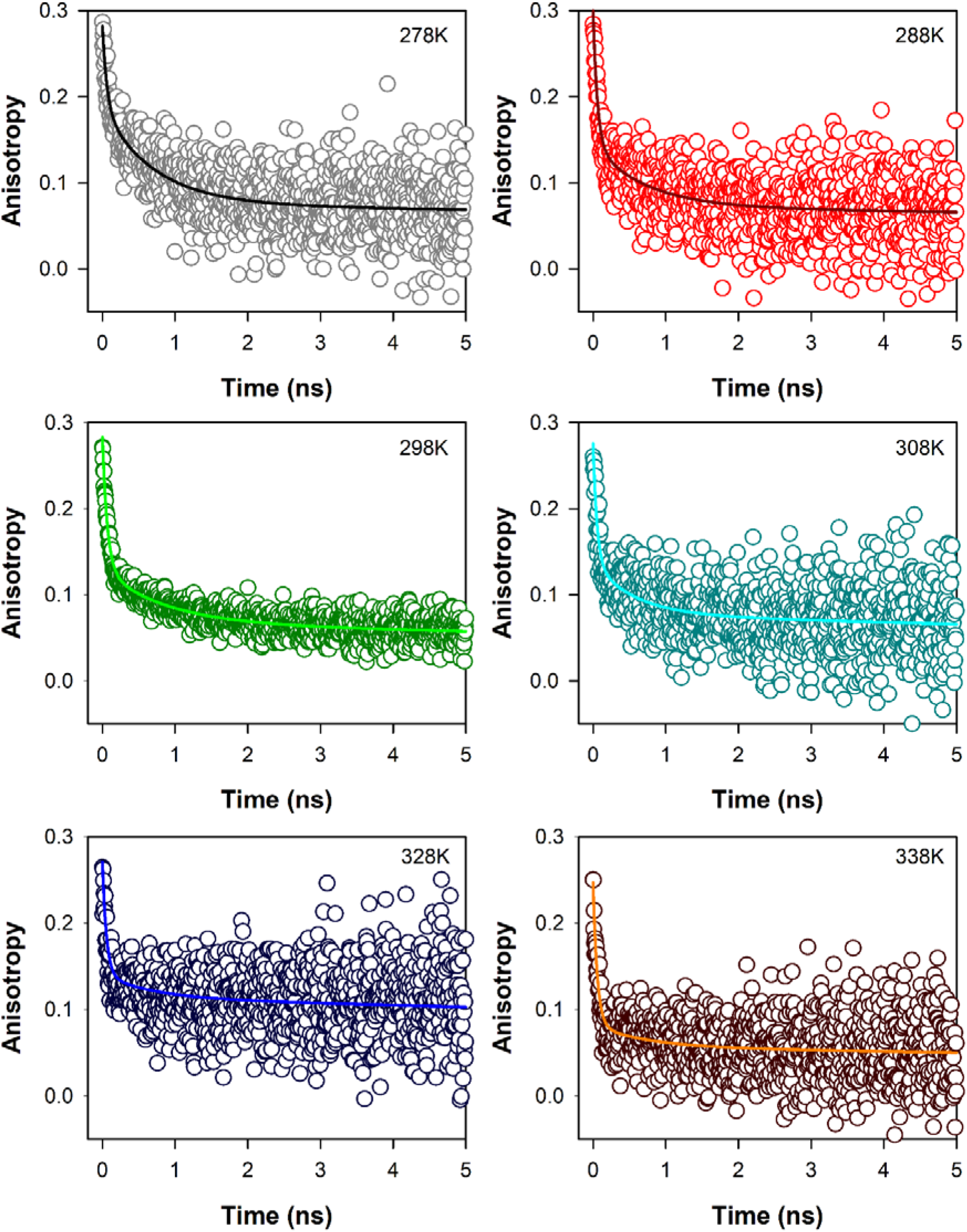
Temperature dependent fluorescence anisotropy of ANS in CHT; solid colored lines represent fits to raw anisotropy data using Eqn 2 as described in the text.

**Table.3:** Time resolved anisotropy of ANS-CHT at different temperatures

| Temperature (K) | $\phi_1$ (ns) | $\phi_2$ (ns) | $\phi_3$ (ns) |
| --- | --- | --- | --- |
| 278 | 0.063 (33 %) | 0.705 (40 %) | 40 |
| 288 | 0.059 (52 %) | 0.707 (24 %) | 40 |
| 298 | 0.067 (54 %) | 0.841 (22 %) | 40 |
| 308 | 0.054 (50 %) | 0.555 (22 %) | 40 |
| 328 | 0.055 (48 %) | 0.676 (10 %) | 40 |
| 338 | 0.057 (67 %) | 0.723 (10 %) | 40 |

**During fitting of anisotropy decay the longer time constant ( $\phi_3$ ) is held fixed at 40 ns**

### Flexibility and mobility analysis

To assess the mobility and flexibility in the different parts of the enzyme root mean square fluctuations (RMSF) of CHT at different temperatures was calculated. RMSF is a measure of the displacement of an amino acid residue around its averaged position during a defined period of time and allows the detection of the regions of high flexibility in a protein. Fig.4(a) shows the RMS fluctuations of CHT at different temperatures. As documented from Fig.4(a) that the loop regions (LRs) experience most of the amino acid residue fluctuations in CHT. There are 5 loop regions (LRs) in particular that form the epicentre of the residue fluctuations: LR1, LR2, LR3, LR4, LR5 (Fig.4b). These LRs comprise of amino acid residues ranging from-LR1 (34-40), LR2 (73-81), LR3 (91-99), LR4 (143-153), LR5 (165-178). All these regions experience fluctuations greater than 2Å, while the other regions experience fluctuations lesser than 2Å with most of them residing at the catalytic S1 pocket. A quick scrutiny of the CHT structure show that the LRs are located around the catalytic pocket the fluctuations of which may control the catalytic activity of CHT. RMS fluctuations however cannot describe the magnitude and direction of the major motions of the protein. Hence normal mode analysis was performed.

**Fig.4:**
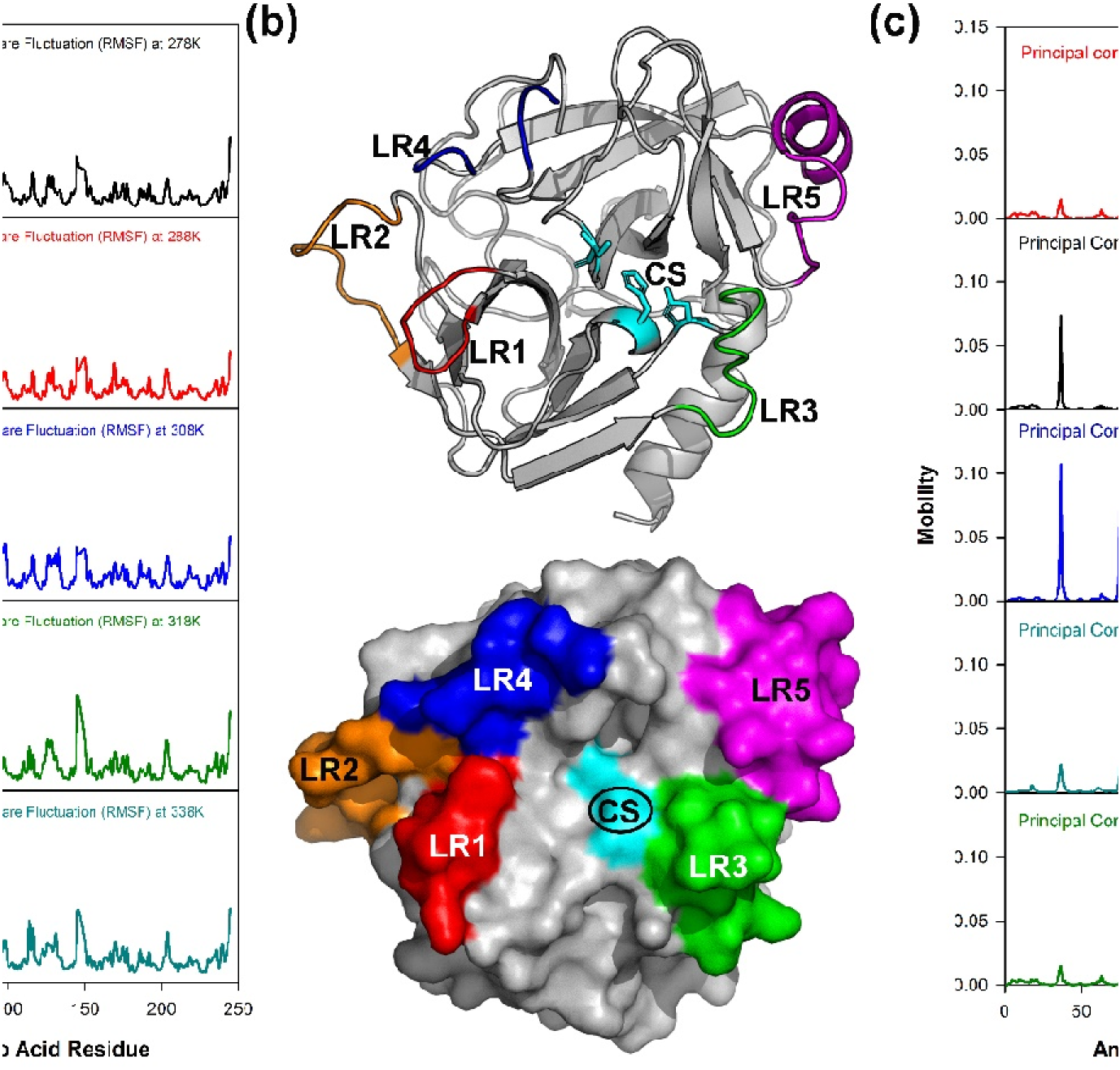
(a)) All atom RMSF of CHT over 30 ns of simulation event at different temperature. The RMSF pattern shows regions of high fluctuation along with regions of low fluctuations. (b) Ribbon representation showing the various loop regions (LRs). The coloured region represents each LRs. LR1 [Red]; LR2 [Orange]; LR3 [Green]; LR4 [Blue]; LR5 [Magenta]. The corresponding amino acid residue for LR1 (34-40), LR2 (73-81), LR3 (91-99), LR4 (143-153), LR5 (165-178). Space filling model showing the positioning of the LRs around the catalytic site (CS); the LRs are represented by same colour scheme. (c) Mobility plot of CHT at different temperatures. The plot shows the epicentre of fluctuations along the amino acid residues across a simulation event. The residues along the LRs are the epicentre of major motions. Regions other than the LRs including the catalytic pocket experience no major fluctuations.

Protein dynamics is demonstrated as a change in molecular conformation as a function of time. To describe the essential motions of the protein over a broad time and spatial scale, protein conformations are best characterized as vector space that that spans a large number of dimensions equal to the number of degrees of freedom (DOF) selected to characterize the motions (33). To generate better insights into the degree of freedom (DOF) and the interpretation of the trajectories principal component analysis (PCA) was performed. The eigenvectors are calculated from the covariance matrix of a simulation trajectory. Along each eigenvector the trajectories are filtered to identify the dominant motion during a simulation event (34, 35). Overall fluctuations of the macromolecules observed from RMSF can often be accounted for by a few low-frequency eigenvectors with large eigenvalues. Study of such eigenvectors and eigenvalues can interprete the essential motions of macromolecules. To study the major motions of CHT all atom principal component analysis (PCA) was applied at different temperatures. Fig.4(c) and Fig.5 illustrates the mobility plot of CHT and the porcupine plot of the eigenvector respectively. The motion of LR1 and LR2 (Fig.4c) clearly shows increase in synchrounous motion with subsequent fall after 308K which correlates with the CHT activity. The porcupine plots (Fig.5) clearly shows the major motions of CHT at different temperatures. Fig.4(c) & Fig.5 specifies the LR regions as the centre of major motions in the protein. The mobility and the porcupine plot specifies the LR regions as the centre of major motions in the protein. At 278K all the LRs experience and act as the centre of major motions. Magnitude of LR motions increase with increase in temperature. At 308K LR1, LR2 and LR5 experience an anti-clockwise motion while LR3 and LR4 moves in clockwise direction. Such synchronous motion can re-orient the LRs positioned around the catalytic pocket making it more accessible to the incoming substrate. Both the porcupine and the mobility plots as well as the motional analysis (ESI Video) indicate the synchronous motion of the LRs with increase in temperature. The presence of sites of synchronous motion could aid in a suitable orientation of the catalytic site for the approaching substrate, thereby, facilitating the enzymatic activity of CHT. However, temperatures above 308K shows the presence of new sites of motion. The mobility plots and the ESI visual inforamtion clearly illustrates new random and independent sites of motion and the loss of synchronous motion in the LRs which may affect catalysis at higher temperatures. [Visual information on the different sites of major motions at different corresponding temperatures are provided as supplementary information].

**Fig.5:**
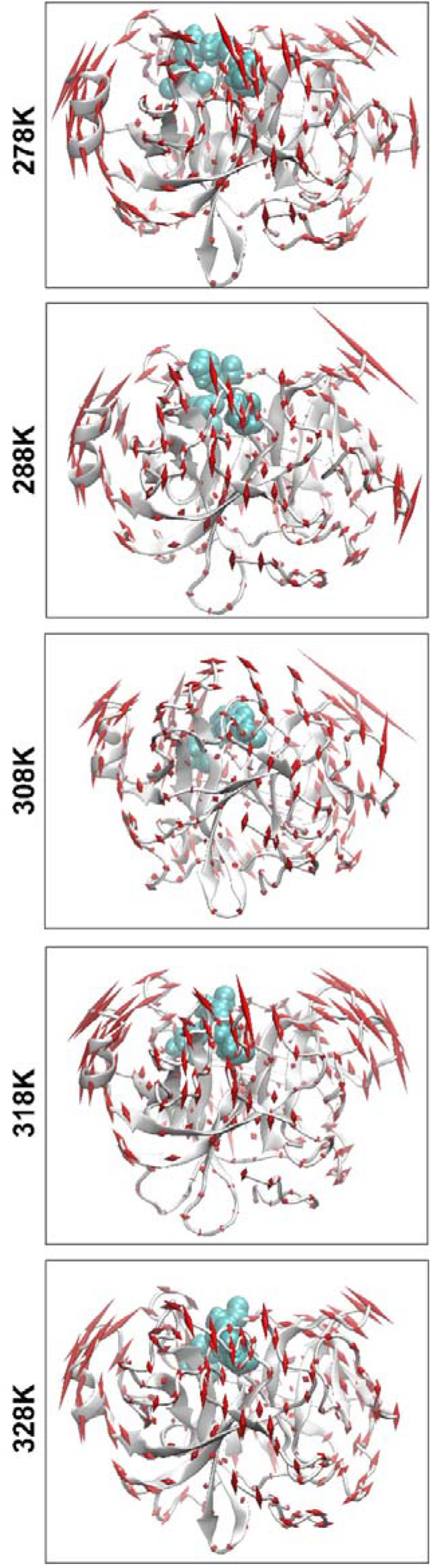
Porcupine plots of CHT at different temperatures. The above model is a backbone trace of CHT. The arrows attached correspond to the direction of each eigenvector while the size of each arrow represents the magnitude of each eigenvalue. The motion of eigenvectors demonstrates synchronous motions of the LRs at 308K. Eigenvector motions become more random and there is loss of concerted eigenvector motion beyond 308K.

### Loop dynamics modulates catalysis in α-chymotrypsin

CHT causes the catalytic hydrolysis of substrate Ala-Ala-Phe-AMC into 7-amido-4-methyl coumarin. The catalytic activity was monitored at different temperatures [278K-338K]. Enhanced product formation suggested the ease of product formation at increased temperatures. Kinetic parameters were calculated from Lineweaver-Burk plot. The catalytic efficiency (k_cat_/K_m_) was a typical bell shaped curve, optimum temperature at 308K (Fig.2) which agrees with a previous study (12). Although a loss in the catalytic efficiency of CHT was noted at 328 and 338K, our thermal stability study at these temperature indicate conformational flexibility as the major architect of the catalytic activity. (Fig.1 & Table. 1). The RMSD and r_gyr_ calculations also corroborate with our thermal stability analysis. Time resolved fluorescence anisotropy of ANS shows significant increase in segmental mobility of the amino acid residues at 308K opposite to the catalytic centre of CHT in comparison to other temperatures below and above 308K. The RMSF calculations show five loop regions (LR1, LR2, LR3, LR4, LR5) as the possible centres of major motions (Fig.4a). A closer inspection of the CHT structure show the LRs are strategically located near the catalytic pocket (Fig.4b). The study further indicates that five LRs may act as possible facilitators of catalytic activity at different temperatures. However, the fluctuations of CHT observed from RMSF cannot provide tangible information regarding the centres of major motions along the simulation trajectory. Principal component analysis (PCA) also indicate the LRs as the epicenter of major motions while the other regions of CHT experience fluctuations of lesser magnitude across all temperatures. The motions of the LRs become less concerted at 278K resulting in reduced catalytic activity (Fig.4c & Fig.5) whereas the concerted motion is increased at 288K reflecting on a change in orientation of the LRs into an open conformation. The concerted synchronous motion of the LRs was maintained at 308K (Fig.4c & Fig.5). The concerted synchronous motion of the LRs was maintained at 308K (Fig.4c & Fig.5) and their open conformation would facilitate an easy access to the incoming substrate leading to increased catalytic efficiency and enhanced turnover at 308K (Table.2). At 338K we observe a loss in concerted motion coupled with the generation of new sites of fluctuations (Fig.4c & Fig.5). This loss in concerted motion of the LRs and amplified randomness may justify the reduction in catalytic efficiency and turnover at temperatures above 308K because increased randomness of the macromolecule and loss of synchronous concerted motion of LRs may lead to a possible dissociation from or inability of the substrate to enter the catalytic pocket.

## Conclusion

Study of the catalytic efficiency of CHT over a range of temperature from 278K to 338K displayed a typical bell shape nature with optimum efficiency being shown at 308K. Secondary structure analysis has indicated maintenance of thermal stability within the studied range of temperature. Polarization-gated fluorescence anisotropy of ANS in CHT display significantly increased segmental mobility of the amino acids residues opposite to the entry of the catalytic centre of the enzyme at 308K in comparison to other temperatures. MD simulations show that synchronous motion of five strategically placed loop regions (LRs) around the catalytic S1 pocket influences and modulates the catalytic activity of CHT at different temperatures. This study manifests the correlation between essential motions in the enzyme to that of its catalytic efficiency which can provide information for future efforts in protein engineering.

## ACKNOWLEDGMENT

SKP thanks the Indian National Academy of Engineering (INAE) for the Abdul Kalam Technology Innovation National Fellowship, INAE/121/AKF.

